# Hydrogen sulfide release via the ACE inhibitor Zofenopril prevents intimal hyperplasia in human vein segments and in a mouse model of carotid artery stenosis

**DOI:** 10.1101/2021.09.13.460108

**Authors:** Diane Macabrey, Céline Deslarzes-Dubuis, Alban Longchamp, Martine Lambelet, Charles K. Ozaki, Jean-Marc Corpataux, Florent Allagnat, Sébastien Déglise

**Author notes:** These authors have equally contributed as first authors (Diane Macabrey, Céline Deslarzes-Dubuis). These authors have equally contributed as senior authors (Florent Allagnat, Sébastien Déglise). Corresponding Author: Florent Allagnat, CHUV-Service de chirurgie vasculaire, Département des Sciences Biomédicales, Bugnon 7A, 1005 Lausanne, Suisse.

## Abstract

**Objectives:** Hypertension is a major risk factor for intimal hyperplasia (IH) and restenosis following vascular and endovascular interventions. Pre-clinical studies suggest that hydrogen sulfide (H_2_S), an endogenous gasotransmitter, limits restenosis. While there is no clinically available pure H_2_S releasing compound, the sulfhydryl-containing angiotensin-converting enzyme inhibitor Zofenopril is a source of H_2_S. Here, we hypothesized that Zofenopril, due to H_2_S release, would be superior to other non-sulfhydryl containing angiotensin converting enzyme inhibitor (ACEi), in reducing intimal hyperplasia.

**Materials:** Spontaneously hypertensive male Cx40 deleted mice (Cx40^−/−^) or WT littermates were randomly treated with Enalapril 20 mg (Mepha Pharma) or Zofenopril 30 mg (Mylan SA). Discarded human vein segments and primary human smooth muscle cells (SMC) were treated with the active compound Enalaprilat or Zofenoprilat.

**Methods:** IH was evaluated in mice 28 days after focal carotid artery stenosis surgery and in human vein segments cultured for 7 days *ex vivo*. Human primary smooth muscle cell (SMC) proliferation and migration were studied *in vitro*.

**Results:** Compared to control animals (intima/media thickness=2.3±0.33), Enalapril reduced IH in Cx40^−/−^ hypertensive mice by 30% (1.7±0.35; p=0.037), while Zofenopril abrogated IH (0.4±0.16; p<.0015 vs. Ctrl and p>0.99 vs. sham-operated Cx40^−/−^ mice). In WT normotensive mice, enalapril had no effect (0.9665±0.2 in control vs 1.140±0.27; p>.99), while Zofenopril also abrogated IH (0.1623±0.07, p<.008 vs. Ctrl and p>0.99 vs. sham-operated WT mice). Zofenoprilat, but not Enalaprilat, also prevented intimal hyperplasia in human veins segments *ex vivo*. The effect of Zofenopril on carotid and SMC correlated with reduced SMC proliferation and migration. Zofenoprilat inhibited the MAPK and mTOR pathways in SMC and human vein segments.

**Conclusion:** Zofenopril provides extra beneficial effects compared to non-sulfhydryl ACEi to reduce SMC proliferation and restenosis, even in normotensive animals. These findings may hold broad clinical implications for patients suffering from vascular occlusive diseases and hypertension.

**Graphical Abstract:** 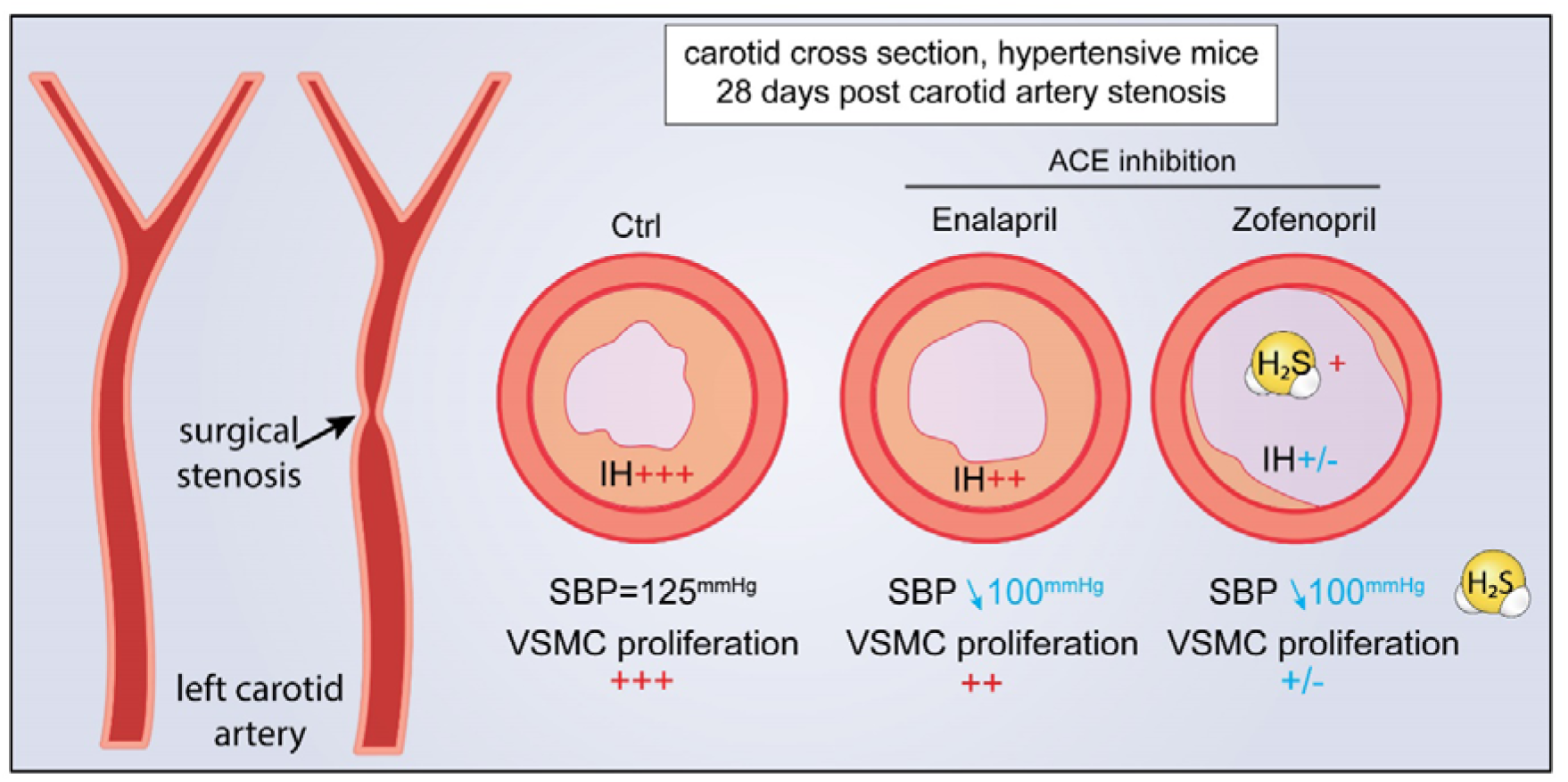

**What this paper adds:** The current strategies to reduce intimal hyperplasia (IH) principally rely on local drug delivery, in endovascular approach. The oral angiotensin converting enzyme inhibitor (ACEi) Zofenopril has additional effects compared to other non-sulfyhydrated ACEi to prevent intimal hyperplasia and restenosis. Given the number of patients treated with ACEi worldwide, these findings call for further prospective clinical trials to test the benefits of sulfhydrated ACEi over classic ACEi for the prevention of restenosis in hypertensive patients.

## INTRODUCTION

Intimal hyperplasia (IH) remains the major cause of restenosis following vascular surgery, leading to potential limb loss and death. IH develops in response to vessel injury, leading to inflammation, vascular smooth muscle cells (SMCs) dedifferentiation, migration, proliferation and secretion of extra-cellular matrix. Despite decades of research, there is no effective medication to prevent restenosis^1^. The only validated therapy against intimal hyperplasia is the local drug-delivery strategy, especially used in endovascular approach. However, this strategy seems to be limited in time^2^, speaking for other complementary oral treatment, targeting either steps involved in IH, such as SMC proliferation, or risk factors for restenosis such as hypertension.

Hydrogen Sulfide (H_2_S) is an endogenously produced gasotransmitter^3^. Pre-clinical studies have shown that H_2_S has cardiovascular protective properties^4^, including reduction of IH^5–7^, possibly via decreased VSMC proliferation^6, 8^. However, there is currently no clinically approved H_2_S donor^9^.

Hypertension is a known risk factor for restenosis and bypass failure^10^. Current guidelines recommend angiotensin converting enzyme inhibitors (ACEi) as first line therapy for the treatment of essential hypertension^11^. Although various ACEi reduce restenosis in rodent models^12^, prospective clinical trials failed to prove efficacy of the ACEi Quinapril^13^ or Cilazapril^14, 15^ for the prevention of restenosis at 6 months following coronary angioplasty. Several *in vitro* studies suggest that the ACEi Zofenopril, due to a sulfhydryl moiety in its structure, releases H_2_S^16–18^. The therapeutic potential of sulfhydryl ACEi Zofenopril has never been tested in the context of restenosis.

The purpose of this study was to test whether Zofenopril, due to its H_2_S-releasing properties, is superior to non-sulfhydryl ACEi in limiting IH in a surgical mouse model of IH *in vivo*, and in an *ex vivo* model of IH in human vein culture. Zofenopril was systematically compared to the non-sulfhydrated ACEi Enalapril.

## MATERIALS and METHODS

### Materials

Drugs and reagents are described in supplemental table S1. Datasets are available at https://doi.org/10.5281/zenodo.5017874

### Experimental group design

All experiments were performed using 8 to 10 weeks old male Cx40-deleted mice (Cx40^−/−^)^19^ and wild-type littermates (WT) mice on a C57BL/6J genetic background. Mice randomly assigned to the experimental groups were treated with the various ACEi at 10mg/Kg/day via the water bottle.

#### Blood pressure experiments

WT (n=22) or Cx40^−/−^ (n=18) mice were randomly divided into 3 groups: control, Enalapril and Zofenopril. Basal systolic blood pressure was measured for 4 days then treatments were initiated and SBP was measured for 10 more days. WT groups were done in parallel (n=22) with Cx40^−/−^ (n=6) untreated mice. Cx40^−/−^ groups (n=18) were done in parallel with WT untreated mice (n=6).

Wild type mice (n=12) were randomly divided into 3 groups: control (n=4), Quinapril (n=4) and Lisinopril (n=4). Basal systolic blood pressure was measured for 4 days then treatments were initiated and SBP was measured for 10 more days.

Systolic blood pressure (SBP) was monitored daily by non-invasive plethysmography tail cuff method (BP-2000, Visitech Systems Inc.) on conscious mice^20^.

#### Mouse carotid artery stenosis model

WT mice (n=26) were divided into 3 groups: Ctrl (n=9), Enalapril (n=9) and zofenopril (n=8). Cx40^−/−^ mice (n=24) were divided into 3 groups: Ctrl (n=11), Enalapril (n=6) and zofenopril (n=7). Seven days post-treatment, intimal hyperplasia was induced via a carotid stenosis.

The carotid artery stenosis (CAS) was performed as previously published^21^. For surgery, mice were anesthetized with Ketamine 80mg/kg and Xylazine 15 mg/kg. The left carotid artery was located and separated from the jugular vein and vagus nerve. Then, a 7.0 PERMA silk (Johnson & Johnson AG, Ethicon, Switzerland) thread was looped under the artery and tightened around the carotid in presence of a 35-gauge needle. The needle was removed, thereby restoring blood flow, albeit leaving a significant stenosis^21^. Buprenorphine 0.05 mg/kg was provided as post-operative analgesic every 12h for 48 hours. Treatment with the ACEi of choice was continued for 28 days post-surgery until organ collection. In another set of surgeries, WT mice (n=17) were randomly divided into 3 groups: control (n=6), quinapril (n=6) and Lisinopril (n=5). Seven days post-treatment, intimal hyperplasia was induced via a carotid stenosis.

All mice were euthanized 28 days post-surgery under general anesthesia by cervical dislocation and exsanguination, perfused with PBS followed by buffered formalin 4% through the left ventricle, and carotids were taken for IH measurements.

All animal experimentation conformed to the *National Research Council: Guide for the Care and Use of Laboratory Animals*^22^. All animal care, surgery, and euthanasia procedures were approved by the Centre Hospitalier Universitaire Vaudois (CHUV) and the Cantonal Veterinary Office (Service de la Consommation et des Affaires Vétérinaires SCAV-EXPANIM, authorization number 3258).

### Ex vivo static human vein culture and SMC culture

Human veins segments were discarded tissue obtained during lower limb bypass surgery. Each native vein was cut into 7mm segments randomly distributed between conditions (D0, D7-Ctrl, D7-enalaprilat or D7-zofenoprilat). One segment (D0) was immediately flash frozen in liquid nitrogen or OCT compound and the other were maintained in culture for 7 days in RPMI-1640 Glutamax supplemented with 10 % FBS and 1% antibiotic solution (10,000 U/mL penicillin G, 10,000 U/mL streptomycin sulphate) at 37°C and 5% CO2, as previously described^6^. The cell culture medium was changed every 48 h with fresh ACEi. 6 different veins/patients were included in this study.

Human vascular smooth muscle cells (VSMC) were prepared and cultured from human saphenous vein segments as previously described^6, 19^. The study protocols for organ collection and use were reviewed and approved by the Centre Hospitalier Universitaire Vaudois (CHUV) and the Cantonal Human Research Ethics Committee (http://www.cer-vd.ch/, no IRB number, Protocol Number 170/02), and are in accordance with the principles outlined in the Declaration of Helsinki of 1975, as revised in 1983 for the use of human tissues. 6 different veins/patients were used in this study to generate VSMC.

### Histomorphometry

Left ligated carotids were isolated and paraffin-embedded. Six 6 μm cross sections were collected every 100 μm and up to 2 mm from the ligature, and stained with Van Gieson Elastic Lamina (VGEL) staining. For intimal and medial thickness, 72 (12 measurements/cross section on six cross sections) measurements were performed^19^. To account for the gradient of IH in relation to the distance from the ligature, the intima thickness was plotted against the distance to calculate the area under the curve of intima thickness. Mean intima and media thickness over the 2mm distance were calculated as well.

For human vein segments, after 7 days in culture, or immediately upon vein isolation (D0), segments were fixed in buffered formalin, embedded in paraffin and cut into 6 μm sections, and stained with VGEL as previously described^6^. For intimal and medial thickness, 96 (4 measurements/photos and 4 photos per cross section on six cross sections) measurements were performed^19^. Two independent researchers blinded to the conditions did the morphometric measurements using the Olympus Stream Start 2.3 software (Olympus, Switzerland)^6, 19^.

### Immunohistochemistry

PCNA (proliferating cell nuclear antigen) immunohistochemistry was performed on paraffin sections as previously described^6^ after antigen retrieval using TRIS-EDTA buffer, pH 9, 17 min in a microwave at 500 watts. Immunostaining was performed using the EnVision +/HRP, DAB+ system according to manufacturer’s instructions (Dako, Switzerland), and counterstained with hematoxylin. One slide per series was assessed and 3 images per section were taken at 200x magnification. Two independent observers unaware of the conditions manually counted the PCNA and hematoxylin positive nuclei.

### Live-cell hydrogen sulfide measurement

Free sulfide was measured in cells using 5 μM SF_7_-AM fluorescent probe as previously described^6^. Fluorescence intensity (λ_ex_ = 495 nm; λ_em_ = 520 nm) was measured continuously in a Synergy Mx fluorescent plate reader (Biotek AG, Switzerland) at 37 °C before and after addition of various compounds, as indicated.

### Persulfidation protocol

Persulfidation protocol was performed using a dimedone-based probe as recently described^23^. Flash-frozen liver was grinded into powder and 20mg of powder was homogenized in 300μl of HEN buffer (100mM HEPES, 1mM EDTA, 100μM neocuproin, 1 vol. % NP-40, 1 wt. % SDS, proteases inhibitors) supplemented with 5mM 4-Chloro-7-nitrobenzofurazan. Proteins were extracted by methanol/chloroform/water protein precipitation and the pellet was resuspended in 200μl of 50mM HEPES-2 wt. % SDS. Protein content was measured using Pierce BCA protein assay kit, and 75μg of proteins were incubated with 25μM final Daz-2-biotin for 1h in the dark at 37°C. Daz-2-biotin was prepared with 1mM Daz-2, 1mM alkynyl biotin, 2mM copper(II)-TBTA, 4mM ascorbic acid with overnight incubation at RT, followed by quenching with 20mM EDTA. Proteins were then extracted by Methanol/Chloroform/Water protein precipitation and the pellets resuspended in 150μl SDS lysis buffer. Protein concentration was measured using the DC protein assay, 10 μg were loaded on SDS-PAGE and the Biotin signal was measured by WB analyses using a streptavidin-HRPO antibody. Protein abundance was normalized to total protein staining using Pierce™ Reversible Protein Stain Kit for PVDF Membranes.

### BrdU staining (VSMC proliferation)

VSMC were grown at 80% confluence on glass coverslips in a 24-well plate and starved overnight in serum-free medium. Then, VSMC were either treated or not (ctrl) with the ACEi of choice for 24h in full medium (RPMI 10% FBS) in presence of 10μM BrdU. All conditions were tested in parallel. All cells were fixed in ice-cold methanol 100% after 24h of incubation and immunostained for BrdU. Images were acquired using a Nikon Eclipse 90i microscope. BrdU-positive nuclei and total DAPI-positive nuclei were automatically detected using the ImageJ software^6^.

### Wound healing assay (VSMC migration)

VSMC were grown at confluence in a 12-well plate and starved overnight in serum-free medium. Then, a scratch wound was created using a sterile p200 pipette tip and medium was changed to full medium (RPMI 10% FBS). Repopulation of the wounded areas was recorded by phase-contrast microscopy over 24 hours in a Nikon Ti2-E live cell microscope. The area of the denuded area was measured at t=0h and t=10h after the wound using the ImageJ software by two independent observers blind to the conditions.

### Western blotting

Human vein segments were washed twice in ice-cold PBS, flash-frozen in liquid nitrogen, grinded to power and resuspended in SDS lysis buffer (62.5 mM TRIS pH6.8, 5% SDS, 10 mM EDTA).

Vascular smooth muscle cells were kept in serum free media overnight. The next morning, complete media was added with the ACEinhibitors. Five hours post-treatment, cells were washed once with ice-cold PBS and directly lysed with Laemmli buffer. Lysates were resolved by SDS-PAGE and transferred to a PVDF membrane (Immobilon-P, Millipore AG, Switzerland). Immunoblot analyses were performed as previously described^6^ using the antibodies described in the **Supplemental Table S1.** Blots were revealed by enhanced chemiluminescence (Immobilon, Millipore) using the ChemiDoc™ XRS+ System and analysed using the Image Lab (BETA2) software, version 3.0.01 (Bio-Rad Laboratories, Switzerland).

### Statistical analyses

All experiments were quantitatively analysed using GraphPad Prism® 8, and results are shown as mean ± SEM. Statistical test details are indicated in the Figure legends.

## RESULTS

### Zofenopril and enalapril similarly lower systolic blood pressure of hypertensive mice

Spontaneously hypertensive Cx40 deleted mice (Cx40^−/−^) and wild-type littermates (WT) were given either 10 mg/kg Zofenopril or 6 mg/kg Enalapril in the drinking water to achieve similar blood lowering effects on hypertensive Cx40^−/−^ mice (**Figure 1A**). Zofenopril also lowered by 6mmHg SBP in normotensive WT mice (**Figure 1B**). The ACEi Enalapril (**Figure 1B**), Quinapril (10 mg/kg) and Lisinopril (10 mg/kg) had no effect on SBP in WT mice (**Figure S1**).

**Figure 1:**
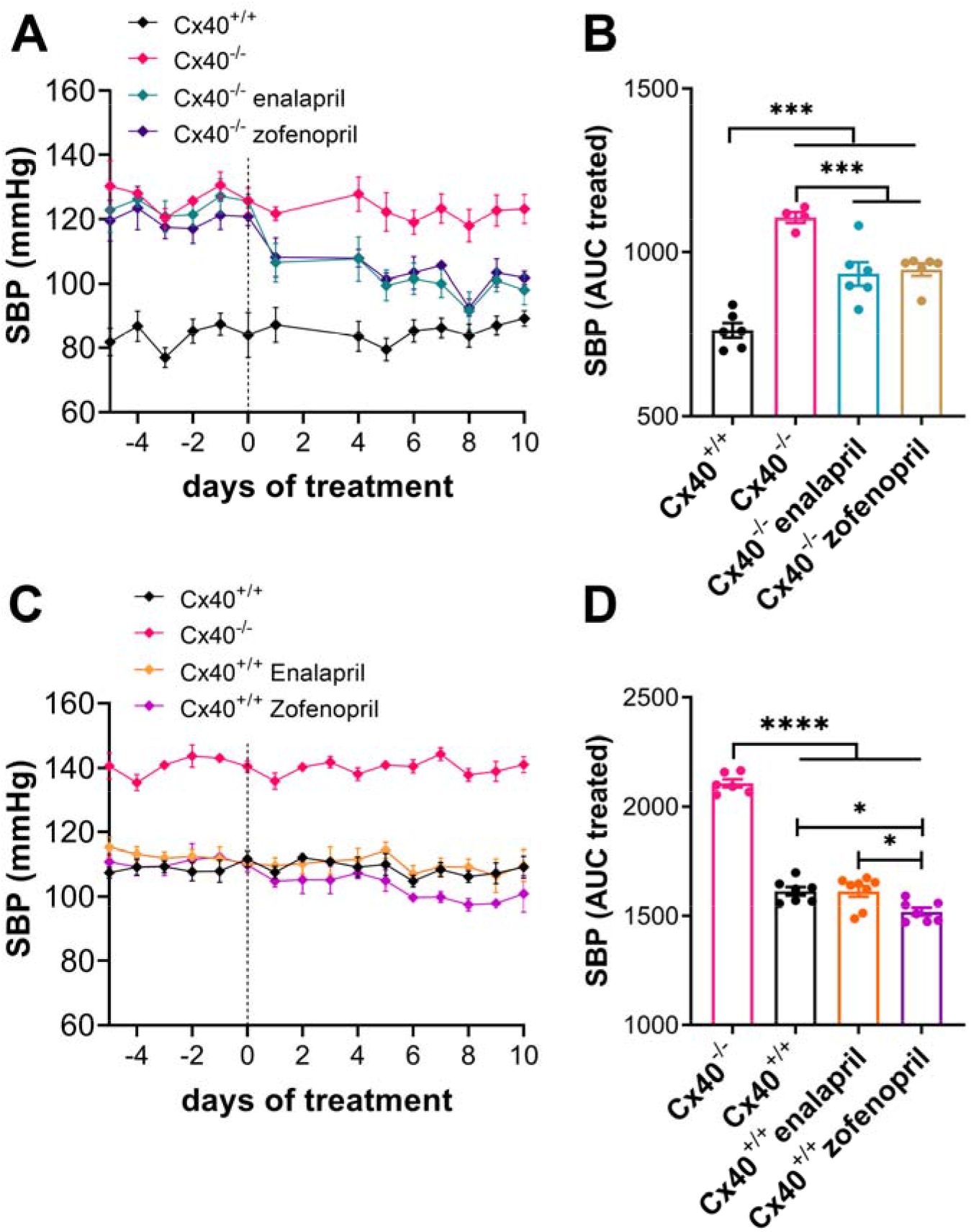
Zofenopril and Enalapril similarly lower systolic blood pressure in hypertensive Cx40^−/−^ mice. **A)** Daily systolic blood pressure (SBP) values (mean±SEM) in WT (n=6) vs. Cx40^−/−^ mice treated or not (n=5) with 10mg/kg Zofenopril (n=6) and 10mg/kg Enalapril (n=6) for the indicated time. **B)** Area under the curve (AUC) of SBP from day 0 to 10. **C)** Daily systolic blood pressure values (mean±SEM) in Cx40^−/−^ (n=5) vs. WT mice treated or not (n=6) with Zofenopril (n=8) and Enalapril (n=8) for the indicated time. D) Area under the curve (AUC) of SBP between day 0 to 10. *p<.05; **p<.01; ***p<.001 as indicated from one-way ANOVA with Tukey’s correction of multiple comparisons.

### Zofenopril is superior to other ACEi in reducing IH in a mouse model of carotid artery stenosis

As expected, the hypertensive mice developed twice more IH than their normotensive littermates following the carotid artery stenosis (CAS) model^21^. Enalapril had a non-specific tendency to reduce IH in Cx40^−/−^ mice (I/M p>.99), while Zofenopril suppressed IH by 90% (I/M p=.0006; **Figure 2, table S2)**. Enalapril had no effect in normotensive WT mice (I/M p>.99), whereas Zofenopril also suppressed IH in those mice (I/M p=.008; **Figure 2, table S3**). The ACEi Quinapril and Lisinopril did not affect IH in WT mice (**Figure S2, table S4**).

**Figure 2.**
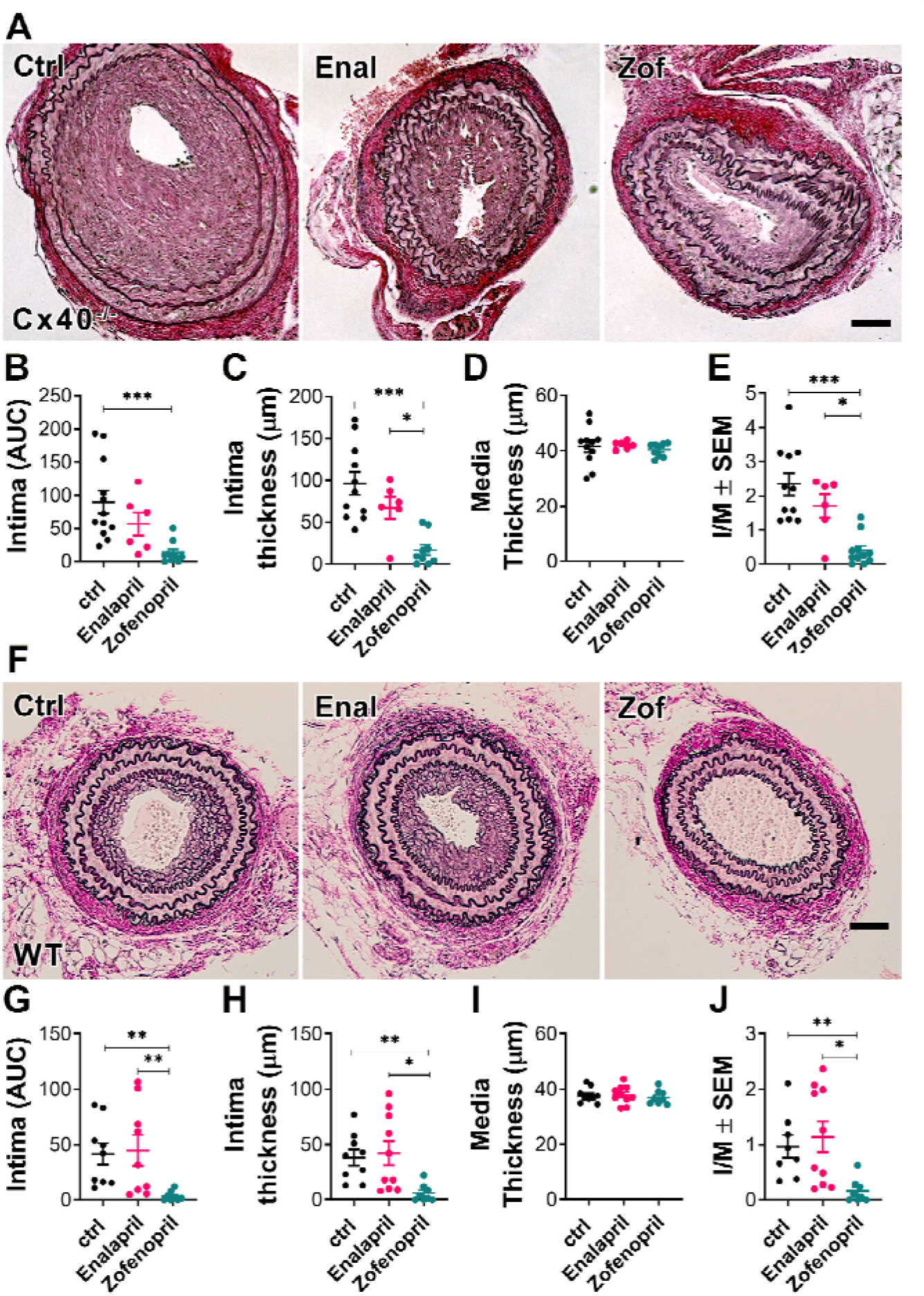
Zofenopril treatment reduces IH in a mouse model of carotid artery stenosis. Cx40^−/−^ (**A-E**) or WT mice (**F-J**), treated or not (Ctrl) with Zofenopril (Zof) and Enalapril (Enal), were submitted to carotid artery stenosis. **A, F**) Representative images of left carotid cross sections stained with VGEL 28 days post-surgery in Cx40−/− (A) or WT (F) mice. Scale bar represents 40 μm. **B-E, G-I**) Morphometric measurements of area under the curve (AUC) of intima thickness (**B, G**) intima thickness (**C, H**), media thickness (**D, I**) and intima over media ratio (**E,J**). Data are presented as scatter plots of 9 to 12 animals per group, with mean±SEM. *p<.05; **p<.01; ***p<.001 as indicated from Kruskal-Wallis test followed by Dunn’s multiple comparisons tests.

### Zofenoprilat prevented the development of IH in human saphenous vein segments

We next tested the effect of Zofenoprilat and Enalaprilat, the active compounds derived from pro-drugs Zofenopril and Enalapril, in our model of IH in *ex vivo* static vein culture^6^. Continuous treatment with 100 μM Zofenoprilat, but not with Enalaprilat, fully blocked the development of IH observed in veins maintained for 7 days in culture in absence of blood flow (D7), compared to initial values in freshly isolated veins (D0) (**Figure 3, table S5**).

**Figure 3:**
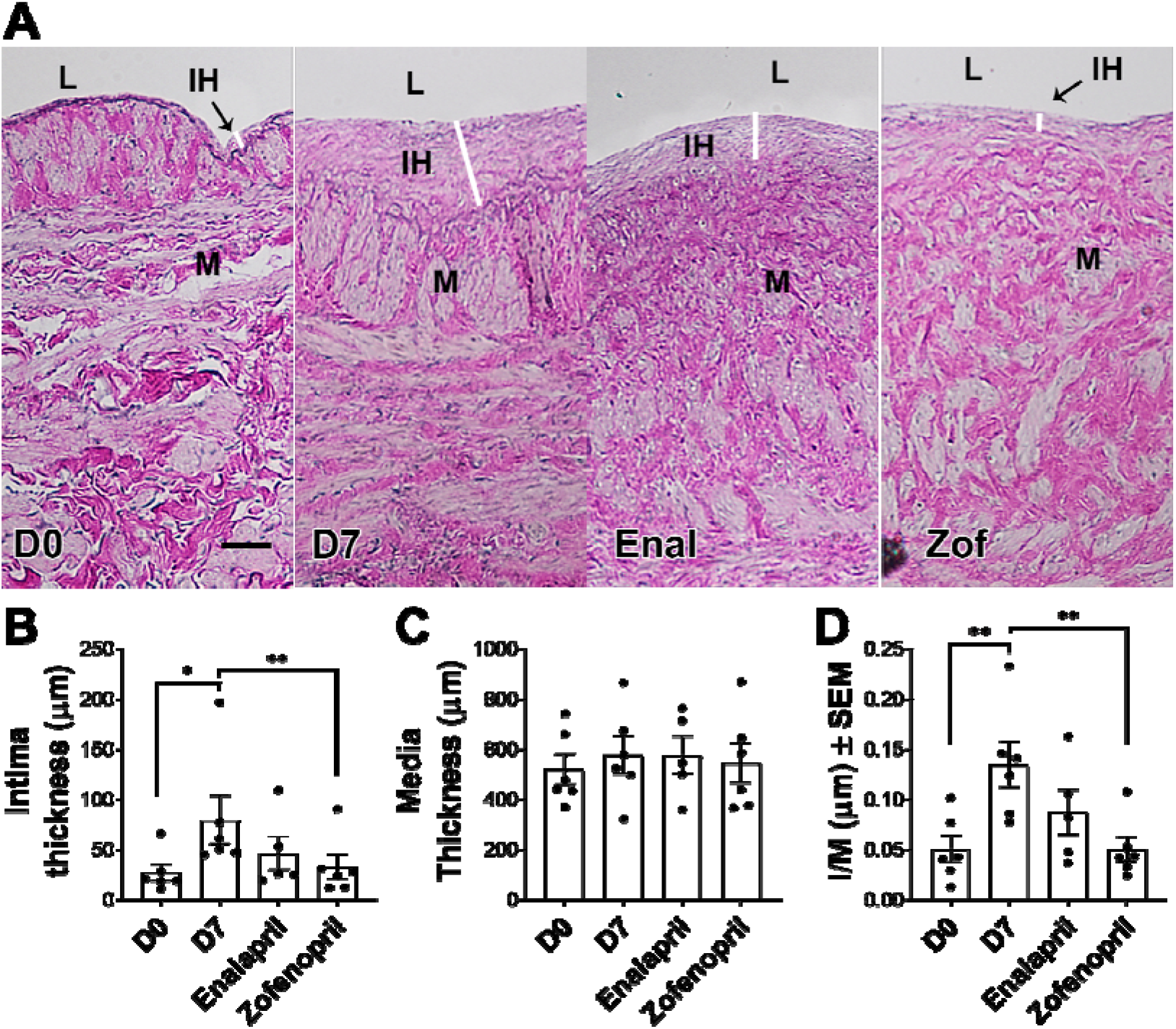
*Ex vivo* treatment with Zofenoprilat prevents the development of IH in human saphenous vein segments. Human great saphenous vein segments obtained from donors who underwent lower limb bypass surgery were put or not (D0, n=6) in static culture for 7 days in presence or not (D7, n=6) of 100μM of Zofenoprilat (n=6) or Enalaprilat (n=5). **A**) Representative VGEL staining. Scale bar= 50μm. B-D) Morphometric measurements of intima thickness (**B**), media thickness (**C**) and intima over media ratio (**D**). Data are presented as scatter plots of 6 different vein/patient with mean±SEM. *p<.05; **p<.01 as indicated from repeated measures one-way ANOVA with post-hoc t-test with Dunnet’s correction of multiple comparisons.

### Zofenoprilat released H_2_S

Besides its ACEi activity, Zofenopril has been proposed to work as an H_2_S donor^16–18^. *In vitro* time-lapse recording of the H_2_S-selective probe SF_7_-AM revealed that Zofenoprilat, but not Enalaprilat, slowly released H_2_S in RPMI medium, compared to the fast-releasing NaHS salt (**Figure 4A**). Similar experiments in presence of live VSMC (**Figure 4B**) confirmed that Zofenoprilat, but not Enalaprilat, increased the SF_7_-AM signal.

**Figure 4.**
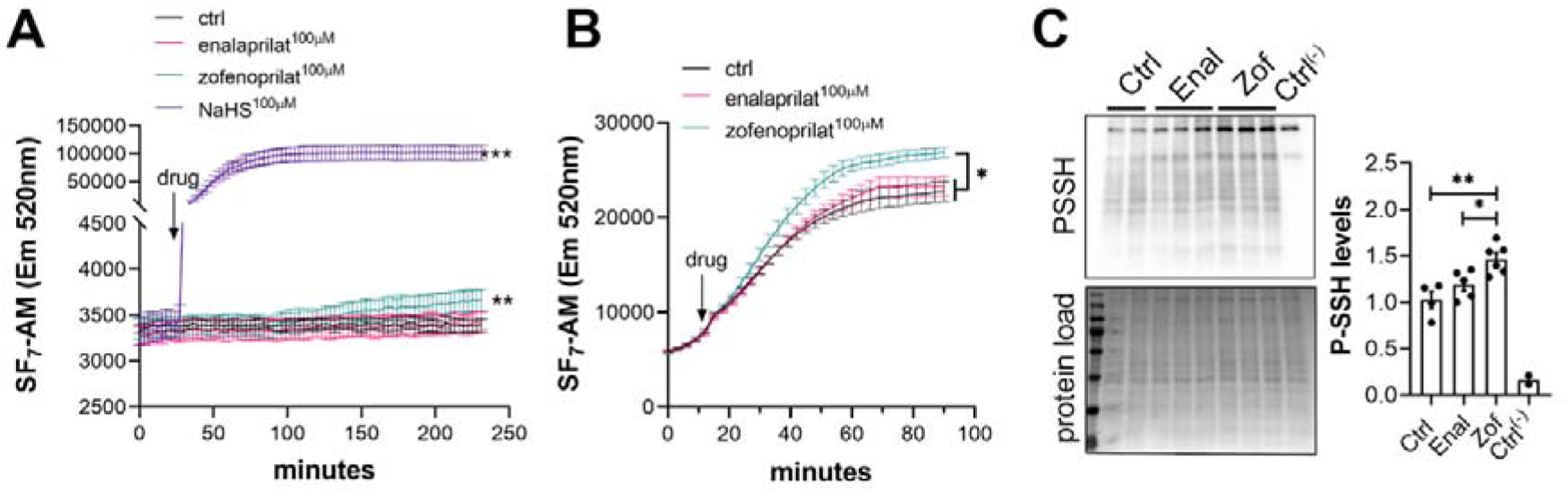
Zofenoprilat release H_2_S as measured by SF_7_-AM fluorescent probe. **A-B)** SF_7_-AM fluorescent signal (mean ± SEM) in a cell free assay in RPMI medium (**A**), in live primary VSMC (**B**) exposed or not (Ctrl) to 100 μM NaHS, 100 μM Zofenoprilat or 100 μM Enalaprilat for the indicated time. *p<.05; **p<.01; ***p<.001 vs. respective Ctrl as determined by repeated measures two-way ANOVA with post-hoc t-test with Tukey’s correction for multiple comparisons. Data are representative of 3 individual experiments. **C)** Global protein persulfidation (PSSH; labelled with DAz-2:Biotin as a switching agent) over total proteins in liver extracts from C57BL/6J male mice treated for two weeks with Enalapril or Zofenopril. Data are presented as scatter plots of 5 to 6 animals/group with mean±SEM with *p<.05, **p<.01, as determined by one-way ANOVA with post-hoc t-test with Tukey’s correction for multiple comparisons.

The biological activity of H_2_S is mediated by post-translational modification of reactive cysteine residues by persulfidation, which modulates protein structure and/or function^9, 23^. We assessed protein persulfidation using a dimedone-based probe as recently described^23^. Zofenopril significantly increased protein persulfidation (PSSH) in liver extracts from mice treated with Enalapril or Zofenopril for two weeks (Figure 4C).

### Zofenopril decreased VSMC proliferation and migration

As various H_2_S donors decrease VSMC proliferation in the context of IH^6, 8^, we next tested the effect of Zofenopril on VSMC. In the CAS model, Zofenopril, but not Enalapril, lowered cell proliferation in the carotid wall as assessed by PCNA staining (**Figure 5A-B**). Zofenoprilat further inhibited the proliferation and migration of primary human VSMC *in vitro*, while Enalaprilat had no effect on proliferation (**Figure 5C-D**) and reduced migration by 20% (**Figure 5E-F**). ACEi Lisinopril and Quinaprilat did not affect VSMC proliferation (**Figure S3**).

**Figure 5.**
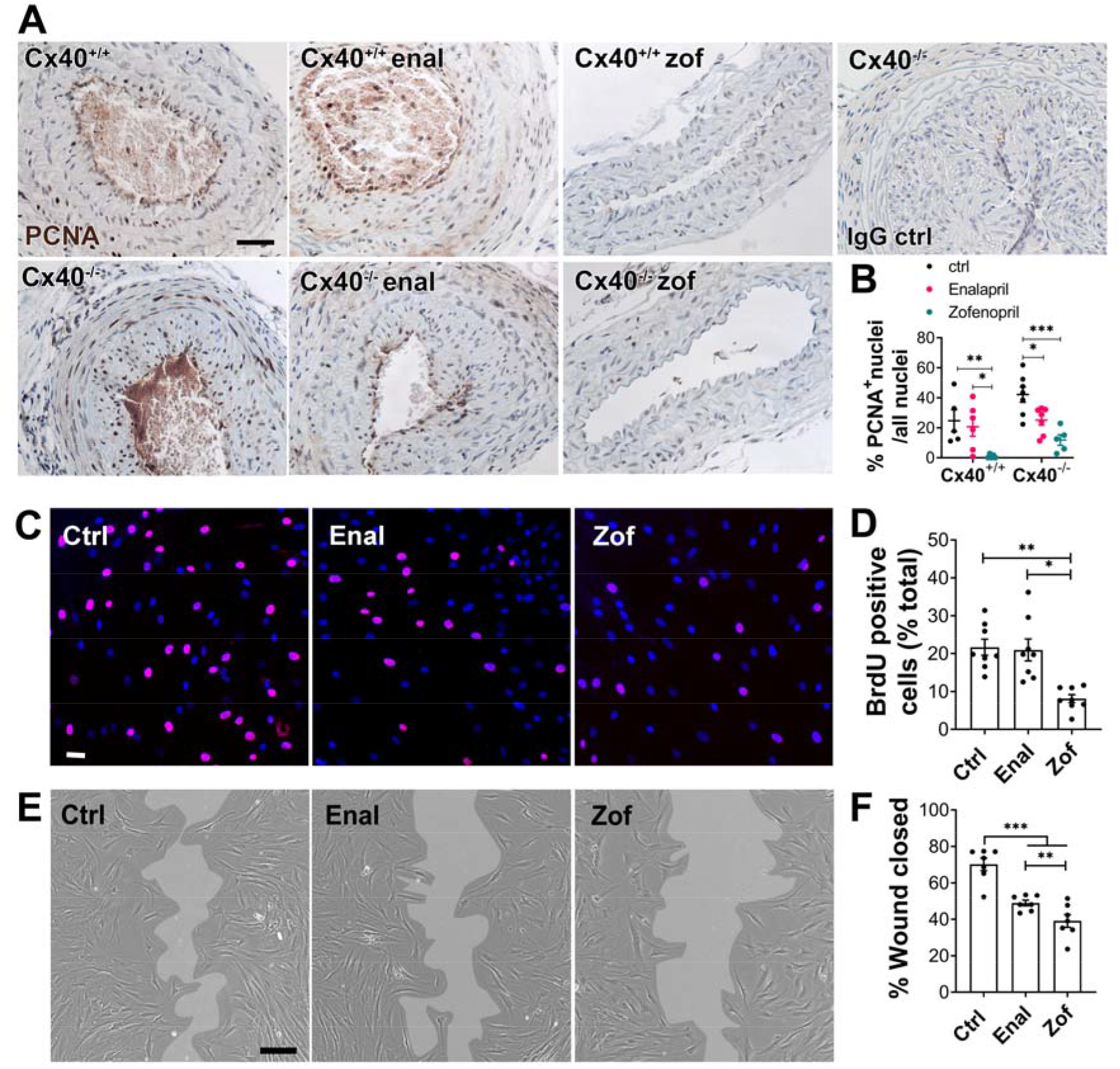
Zofenopril treatment reduces cell proliferation in a mouse model of carotid artery stenosis. **A-B)** PCNA immunostaining 28 days post carotid artery stenosis in WT (Cx40^+/+^) or Cx40^−/−^ mice treated or not (Ctrl) with Zofenopril (Zof) and Enalapril (Enal). **A**) Representative images of PCNA-positive nuclei (brown) and negative nuclei (haematoxylin-stained blue nuclei). Scale bar represents 40 μm. **B**) Quantitative assessment of PCNA positive cells over total cells of 5 to 8 animals/group with mean±SEM. *p<.05; **p<.01; ***p<.001, as determined by two-way ANOVA with post-hoc t-test with Sidak’s correction of multiple comparisons. **C-D**) Primary human vascular smooth muscle cells (VSMC) were exposed or not (Ctrl) to 100 μM Zofenoprilat or Enalaprilat for 24 h in presence of BrdU. **C**) Representative images of BrdU-positive nuclei (pink) and DAPI-stained nuclei (blue). Bar scale represents 10 μm. **D**) Proliferation was calculated as the percentage of BrdU-positive nuclei over total nuclei. Data are scatter plots of 8 independent experiments with mean±SEM with *p<.05, **p<.01, as determined by repeated measures one-way ANOVA with post-hoc t-test with Dunnet’s correction for multiple comparisons. **E-F**) Wound healing assay with VSMC exposed or not (Ctrl) to 100 μM Zofenoprilat or Enalaprilat. **E**) Representative images of VSMC in brightfield 10 hours post wound. Scale bar represents 50 μm. **F**) Data are scatter plots of 7 independent experiments with mean±SEM of wound area after 10h, expressed as a percentage of the initial wound area. **p<.01, ***p<.001, as determined by repeated measures one-way ANOVA with post-hoc t-test with Dunnet’s correction for multiple comparisons.

### Zofenoprilat inhibited the MAPK and mTOR pathways

The MAPK and mTOR signalling pathways contribute to VSMC proliferation in the context of IH^24^. Western blot analyses revealed that Zofenoprilat reduced by 50% the levels of P-ERK1,2, P-p38 and P-S6RP in cultured VSMC, while Enalaprilat had no effect (**Figure 6A-F**). Moreover, P-S6RP and P-ERK1,2 levels were also decreased by Zofenoprilat in human vein segments placed in culture for 7 days (**Figure 6G-I**).

**Figure 6:**
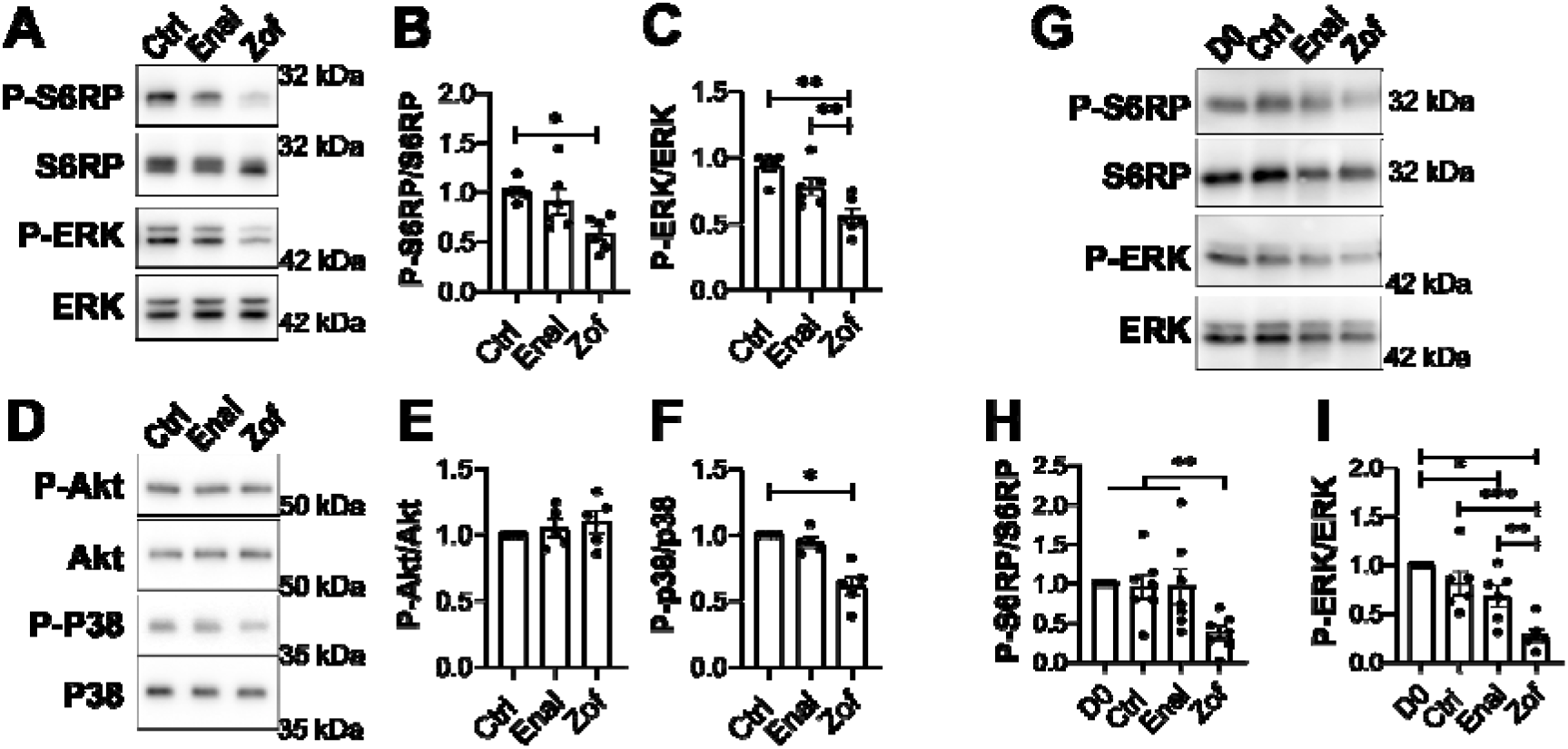
Zofenoprilat inhibits ERK and S6RP phosphorylation. **A-F**) Western Blot analyses from VSMC exposed or not (Ctrl) to 100 μM Zofenoprilat or Enalaprilat for 5h. **A, D**) Representative Western Blot for P-S6RP and total S6RP, P-ERK and total ERK, P-Akt and total Akt, P-p38 and total p38. **B-C; E-F**) Quantitative assessment of 6 independent experiments, normalized to their respective Ctrl condition, with mean±SEM. *p<.05, **p<.01, as determined by repeated measures one-way ANOVA with post-hoc t-test with Dunnet’s correction for multiple comparisons. **G-I**) Western Blot analyses from human vein segments exposed or not (Ctrl) to 100 μM Zofenoprilat or Enalaprilat for 7 days. **G**) Representative Western Blot for P-S6RP and total S6RP, PERK and total ERK. **H-I**) Quantitative assessment of 7 different veins, normalized to their respective Ctrl condition, with mean±SEM. *p<.05, **p<.01, ***p<.001, as determined by repeated measures one-way ANOVA with post-hoc t-test with Dunnet’s correction for multiple comparisons.

## DISCUSSION

In this study, we hypothesized that Zofenopril, an ACEi with a free thiol moiety acting as an H_2_S donor, would be more efficient than other ACEi to inhibit IH in the context of hypertension. Not only Zofenopril is more potent than Enalapril in reducing IH in hypertensive Cx40^−/−^ mice, it also suppresses IH in normotensive condition, where other ACEi have no effect. Furthermore, Zofenopril prevents IH in human saphenous vein segments in absence of blood flow. The effect of Zofenopril on IH correlates with reduced VSMC proliferation and migration and decreased activity of the MAPK and mTOR pathways.

Several pre-clinical studies have shown that that SBP-lowering medication such as ACEi reduce IH^12^, which prompt the large-scale MERCATOR/MARCATOR^14, 15^ and PARIS clinical trials^13^. Here, we also observed that lowering SBP using the ACEi Enalapril had a non-significant tendency to protect from IH in hypertensive mice. However, Enalapril, Quinapril and Lisinopril, had no effect in normotensive WT mice. The fact that the sulhydrated ACEi Zofenopril almost abrogated IH in hypertensive and normotensive mice strongly supports that this ACEi provides additional effects independent of its ACEi activity, as previously suggested^16–18^. Of interest, the SMILE clinical trials concluded that, compared to placebo or Ramipril, Zofenopril reduces the 1-year risk of cardiovascular events after acute myocardial infarction^25^. These benefits might be related to H_2_S release by Zofenopril, as pre-clinical studies consistently show that H_2_S supplementation promote recovery after after acute myocardial infarction^4^.

Zofenopril has been proposed to work as a H_2_S donor in several studies^16–18^. Here, we confirmed that Zofenoprilat releases detectable amounts of H_2_S. H_2_S modifies proteins by post-translational persulfidation (S-sulfhydration) of reactive cysteine residues, which modulate protein structure and/or function^23^. Here, we further observed that Zofenopril increases overall protein persulfidation *in vivo*, suggesting that Zofenopril generates H_2_S *in vivo* as well.

We and others previously demonstrated that various H_2_S donors inhibit VSMC proliferation^6, 8, 26^. Consistently, we confirmed that Zofenopril inhibits VSMC proliferation and migration *in vitro* and reduces cell proliferation in the carotid wall *in vivo*. Although the exact mecanisms of action of Zofenoprilat and H_2_S remain to be elucidated, we demonstrated that Zofenoprilat inhibits the MAPK and mTOR signalling pathways, which contribute to VSMC proliferation and neointima formation^24^. Overall, our data strongly suggest that Zofenopril acts similarly to other known H_2_S donors to limit IH through inhibition of the MAPK and mTOR signalling pathways, leading to decreased VSMC proliferation and migration.

Overall, our data suggests that Zofenopril might show benefits against restenosis in patients unlike other ACEi. These findings raise the question as to whether the scientific community was not too quick to discard the whole class of ACEi as a treatment of restenosis based on the disappointing results of the MERCATOR/MARCATOR^14, 15^ and PARIS trials^13^. In the last decade, many efforts have been put on the development of local drug delivery strategy, well adapted to endovascular interventions. However, this strategy seems to bring great improvement in the mid-term but not in the long-term follow-up^2^. Thus, a more chronic approach, sustaining the early effect on cell proliferation and IH inhibition, should be encouraged. Such a strategy relies on oral medication, which is also better adapted to open surgery.

### Nevertheless, our study carries some limitations

Firstly, numerous oral drugs have been clincially tested over the years to limit restenosis and, in most trials, the pharmacologic treatment of restenosis failed to show positive results, despite promising results obtained in experimental models^27^. While there is no doubt that pre-clinical models have significantly advanced our understanding of the mechanisms of restenosis formation, none of them fully mimic restenosis in human. The genetic model of renin-dependent hypertension used in that study is also rarely observed in patients, which have complex multifactorial essential hypertension. Additional studies reflecting better the patients’comorbidities (dyslipidemia, renal insufficiecy, smoking, atherosclerosis, etc.) with a vein bypass model and larger animal models, or small phase II clinical trial, are required before testing the benefits of Zofenotpril in a large, phase III clinical trials.

Secondly, although Zofenopril was the only ACEi providing benefits in normotensive condition, we cannot exclude that other ACEi not tested here could work as well. We further acknowledge that pharmacokinetics and pharmacodynamics differences between Zofenopril and other ACEi may contribute to the superiority of Zofenopril. Zofenopril is more lipophilic and may have better tissue penetration than Enalapril or Ramipril, which may have an impact beyond the effect of H_2_S liberated by Zofenopril. However, it has been shown that vessel wall penetration of various ACEi is independent of lipophilia and that the endothelium constitutes no specific barrier for the passage of ACE inhibitors^28^.

Finally, our working hypothesis is that Zofenopril inhibits VSMC proliferation via direct release of H_2_S at the level of the media of vessel. However, we could not ascertain that H_2_S is released at the level of the VSMC. H_2_S^9^ and Zofenoprilat^17, 18^ have been shown to promote endothelial cell function, including proliferation and migration. Thus, we cannot exclude that Zofenopril limits IH via a positive effect on endothelial cells. Further studies are required to carefully assess the impact of Zofenopril on the endothelium and quantify H_2_S in vascular tissue.

## Conclusion

Under the conditions of these experiments, Zofenopril is superior to Enalapril in reducing IH and provides beneficial effect against IH in mice and in a model of IH in human vein segments *ex vivo*. Our data strongly support that Zofenopril limits the development of IH via H_2_S release, independently of its ACEi activity. The effects of Zofenopril correlate with reduced MAPK and mTOR pathways activities, leading to decreased VSMC proliferation and migration.

Given the number of patients treated with ACEi worldwide, these findings may have broad implications for the treatment of patients suffering from peripheral atherosclerotic disease undergoing revascularization, and beyond. Our results warrant further research to evaluate the benefits of Zofenopril in limiting restenosis and eventually prospective clinical trials to test the superiority of sulfhydrated ACEi on restenosis over other ACEi.

## Abbreviations

ACE: angiotensin converting enzyme
CAS: carotid artery stenosis
Cx40: connexin 40
H_2_S: hydrogen sulfide
IH: intimal hyperplasia
PCNA: proliferating cell nuclear antigen
SBP: systolic blood pressure
VSMC: vascular smooth muscle cells
VGEL: Van Gieson elastic lamina

## Acknowledgements

We thank Prof. Jacques-Antoine Haefliger for giving us the Cx40^−/−^ mice. We thank the mouse pathology facility for their services in histology (https://www.unil.ch/mpf).

