## Supplemental data and methods for "Hydrogen sulfide release via the ACE inhibitor Zofenopril prevents intimal hyperplasia in human vein segments and in a mouse model of carotid artery stenosis"

DATA SUPPLEMENT

**Supplemental Table S1: Materials**

| **product** | **Vendor** | **Catalog #** | **Working concentration** |
| --- | --- | --- | --- |
| **Zofenopril 30mg** | Mylan SA |  | 10mg/kg/day |
| **Enalapril 20mg** | Mepha Pharma |  | 6mg/kg/day |
| **Accupro “Quinaprilum” 20mg (Quinapril)** | Pfizer |  | 10mg/kg/day |
| **Lisinopril 10mg** | Mepha Pharma |  | 10mg/kg/day |
| **Zofenoprilat** | Santa Cruz Biotech | Sc-220412 | 100µM |
| **Enalaprilat** | Sigma-Aldrich | e9858 | 100µM |
| **Quinaprilat** | Santa Cruz Biotech | sc-208193A | 100µM |
| **Lisinopril (pure active compound)** | Santa Cruz Biotech | sc-205378 | 100µM |
| **P-S6RP (Ser 236)** | Cell Signaling Technology | #4858 | 1/1000 |
| **S6RP** | Cell Signaling Technology | #2217 | 1/2000 |
| **P-ERK1,2 (Tyr 202-204)** | Cell Signaling Technology | #4370 | 1/2000 |
| **ERK1,2** | Cell Signaling Technology | #4695 | 1/5000 |
| **P-p38 (Thr 180/182)** | Cell Signaling Technology | #9211 | 1/1000 |
| **p38** | Cell Signaling Technology | #9212 | 1/1000 |
| **P-AKT (Ser 473)** | Cell Signaling Technology | #4051 | 1/1000 |
| **AKT** | Cell Signaling Technology | #9272 | 1/1000 |
| **HRP-Streptavidin** | Sigma-Aldrich | RABHRP3 | 1/5000 |
| **Anti-Rabbit HRP** | Invitrogen | #31460 | 1/20000 |
| **Anti-mouse HRP** | Jackson Immunoresearch | 115-035-146 | 1/15000 |
| **BrdU** | BD Biosciences | 555627 | 1/200 |
| **PCNA** | Dako, Baar, Switzerland | M0879 | 1/100 |
| **SF7-AM fluorescent probe** | Sigma-Aldrich | 748110 |  |
| **Daz-2** | Cayman Chemicals | 13382 |  |
| **alkynyl biotin** | Cayman Chemicals | 13038 |  |
| **copper(II)-TBTA** | Lumiprobe | 21050 |  |
| **4-Chloro-7-nitrobenzofurazan** | Sigma-Aldrich | 163260 |  |
| **Pierce™ BCA Protein Assay Kit** | ThermoFisher Scientific | 23225 |  |
| **Pierce™ Reversible Protein Stain Kit for PVDF Membranes** | Thermo Fisher Scientific | 24585 |  |
| **DC™ Protein Assay Kit I** | Bio-Rad | 5000111 |  |
| **Dako EnVision® + Dual Link System- HRP (DAB+)** | Agilent | K4065 |  |
| **EGM™-2 (Endothelial Cell Growth Medium-2 BulletKit™)** | Lonza | CC-3162 |  |
| **RPMI 1640 Medium, GlutaMAX™ Supplement** | Thermo Fisher Scientific | 61870036 |  |
| **Ketasol-100 (Ketamine)** | Gräub E.Dr.AG, Switzerland | QN01AX03 |  |
| **Rompun® 2% (Xylazine)** | Provet AG, Switzerland | QN05CM92 |  |
| **Temgesic® (Buprenorphin)** | Reckitt Benckiser | N02AE01 |  |

**Supplemental Table S2: Detailed results table for Cx40^-/-^ mice**

| **Cx40^-/-^ mice** | | | |
| --- | --- | --- | --- |
|  | Ctrl | Enalapril | Zofenopril |
| **Intima Thickness** | 96.1±13.7 (n=11) | 67.1±13.3 (n=6) | 16.8±6.3 (n=9) |
| **Media Thickness** | 41.7±2.1 (n=11) | 42.1±0.6 (n=6) | 40.4±0.8 (n=9) |
| **I/M** | 2.34±0.33 (n=11) | 1.7±0.35 (n=6) | 0.39±0.13 (n=9) |

**Supplemental Table S3: Detailed results table for WT mice**

| **Wild type mice** | | | |
| --- | --- | --- | --- |
|  | Ctrl | Enalapril | Zofenopril |
| **Intima Thickness** | 38.3±7.2 (n=9) | 42.1±10.9 (n=10) | 6±2.8 (n=8) |
| **Media Thickness** | 37.7±0.9 (n=9) | 37.9±1.2 (n=10) | 36.9±0.9 (n=8) |
| **I/M** | 0.96±0.21 (n=9) | 1.14±0.28 (n=10) | 0.16±0.07 (n=8) |

**Supplemental Table S4: Detailed results table for WT mice**

| **Wild type mice** | | | |
| --- | --- | --- | --- |
|  | Ctrl | Quinapril | Lisinopril |
| **Intima Thickness** | 33.4±7.1 (n=6) | 40.5±9.1 (n=6) | 37.7±12.4 n(5) |
| **Media Thickness** | 34.8±1.2 (n=6) | 34.5±1.4 (n=6) | 36.6±1.4 (n=5) |
| **I/M** | 1.06±0.25 (n=6) | 1.27±0.38 (n=6) | 1.08±0.37 (n=5) |

**Supplemental Table S5: Detailed results table for human vein segments**

| **Human vein segments** | | | | |
| --- | --- | --- | --- | --- |
|  | D0 | D7 | D7-Enalapril | D7-Zofenopril |
| **Intima Thickness** | 27.6±8.1 (n=6) | 80.1±23.9 (n=6) | 47.1±16.7 (n=5) | 33.4±11.9 (n=6) |
| **Media Thickness** | 522.3±59.7 (n=6) | 578.9±74.6 (n=6) | 578.4±73.5 (n=5) | 548.7±79.1 (n=6) |
| **I/M** | 0.05±0.01 (n=6) | 0.13±0.02 (n=6) | 0.08±0.02 (n=5) | 0.05±0.01 (n=6) |

### SUPPLEMENTAL FIGURES

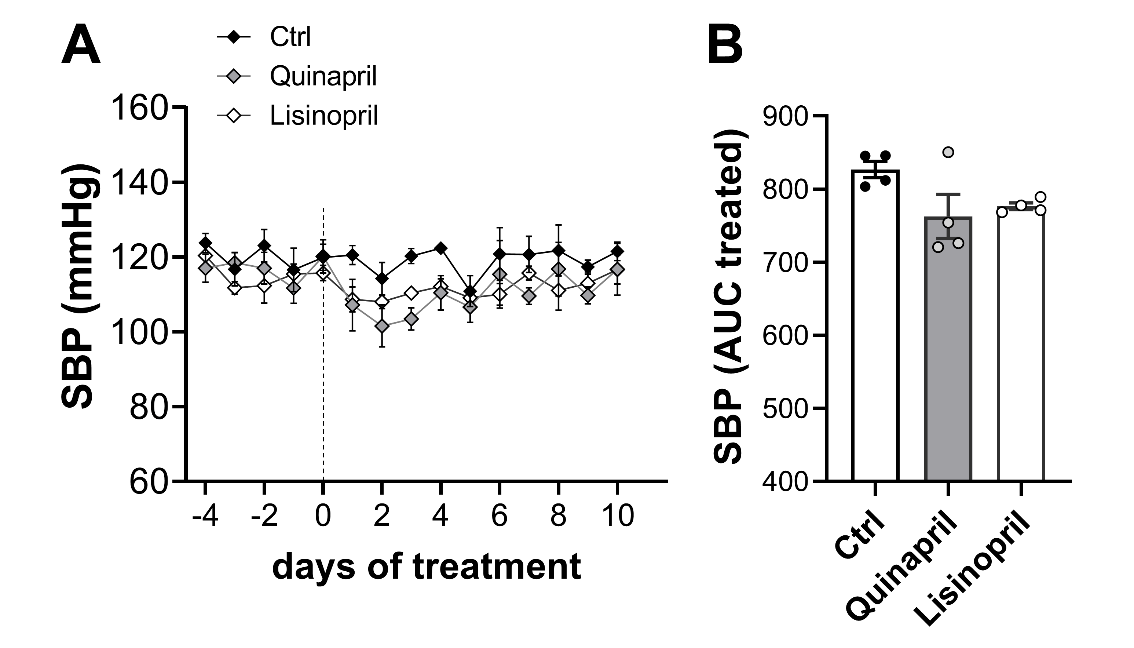

**Figure S1. Quinapril and Lisinopril do not reduce SBP in WT mice**

**A)** Daily systolic blood pressure (SBP) values (mean±SEM) in WT mice treated or not (Ctrl) with Quinapril (10 mg/kg) or Lisinopril (10 mg/kg) in 4 animals per group with mean±SEM. **B)** area under the curve (AUC) of SBP from day 0 to 10. No statistical difference as determined by one-way ANOVA with Tukey’s correction for multiple comparisons.

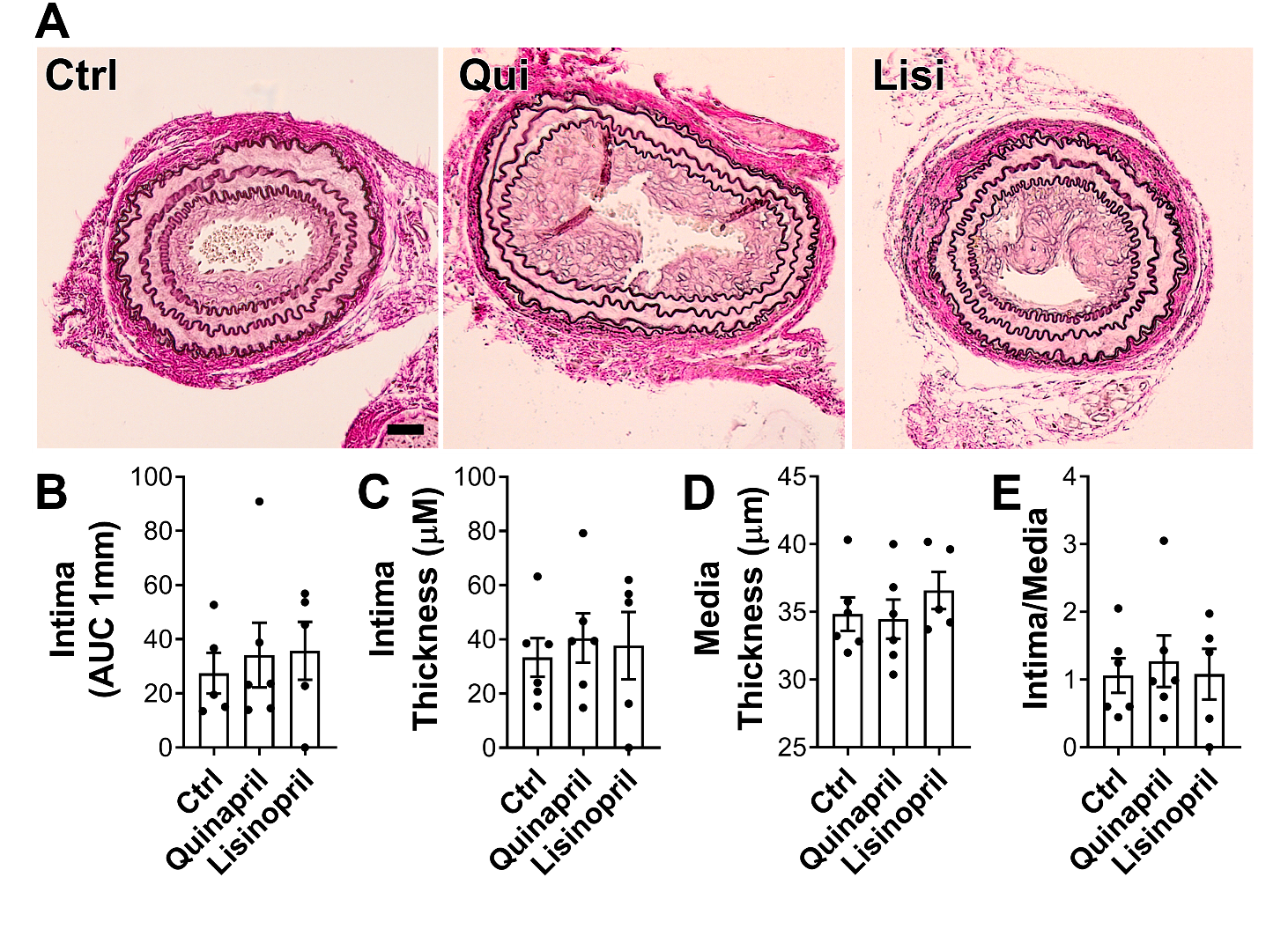

### Figure S2. Quinapril and Lisinipril do not reduce IH in a mouse model of carotid artery stenosis

WT mice treated or not (Ctrl) with Quinapril (Qui) or Lisinopril (Lisi), were submitted to carotid artery stenosis. **A**) Representative left carotid cross sections stained with VGEL 28 days post-surgery. Scale bar represents 30 µm. **B-D**) Data are morphometric measurements of area under the curve (AUC) of intima thickness (**B**) intima thickness **(C**), media thickness (**D**) and intima over media ratio (**E**). Data are scatter plots of 5 to 6 animals per group with mean±SEM. No statistical difference as calculated by two-way ANOVA with Tukey’s correction of multiple comparisons.

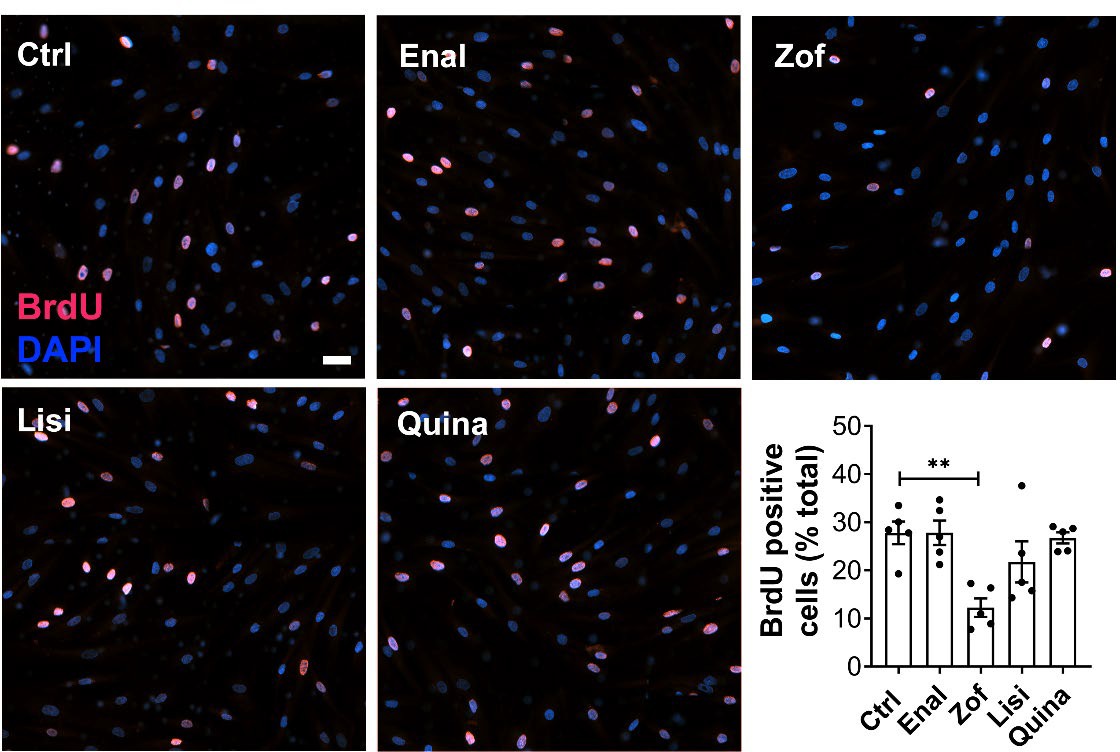

### Figure S3. Quinaprilat and Lisinopril do not reduce VSMC proliferation.

Primary human vascular smooth muscle cells (VSMC) were exposed or not (Ctrl) to 100 µM Zofenoprilat, Enalaprilat, Lisinopril or Quinaprilat for 24 h in presence of BrdU. Bar scale represents 20 µm. Proliferation was calculated as the percentage of BrdU-positive nuclei (pink) over total DAPI-stained nuclei (blue). Data are shown as scatter plots of 5 independent experiments with mean±SEM. **p<.01 as determined by paired one-way ANOVA with post-hoc t-test with Dunnet’s correction for multiple comparisons.
